# Decoupling of the Onset of Anharmonicity between a Protein and Its Surface Water around 200 K

**DOI:** 10.1101/2023.10.18.562890

**Authors:** Lirong Zheng, Bingxin Zhou, Banghao Wu, Yang Tan, Juan Huang, Madhusudan Tyagi, Victoria García Sakai, Takeshi Yamada, Hugh O’Neill, Qiu Zhang, Liang Hong

## Abstract

The protein dynamical transition at ∼ 200 K, where the biomolecule transforms from a harmonic, non-functional form to an anharmonic, functional state, has been thought to be slaved to the thermal activation of dynamics in its surface hydration water. Here, by selectively probing the dynamics of protein and hydration water using elastic neutron scattering and isotopic labelling, we found that the onset of anharmonicity in the two components around 200 K are decoupled. The one in protein is an intrinsic transition, whose characteristic temperature is independent of the instrumental resolution time, but varies with the biomolecular structure and the amount of hydration, while the one of water is merely a resolution effect.

## Introduction

It is well established that the internal dynamics of a protein is crucial for its functions, including allosteric conformational changes (1), ligand binding (2) and enzymatic reactions (3). In particular, hydrated proteins exhibit a dynamical transition around 200 K, across which the slope of the temperature dependence of the atomic displacements changes significantly and the biomolecule transforms from a rigid, harmonic state, to a flexible, anharmonic form (4-12). Although exceptions have been reported (50), the dynamical transition has been linked to the thermal onset of function in a number of proteins, e.g, myoglobin (13), ribonuclease (6), elastase (14) and bacteriorhodopsin (15), all of which become inactive below the dynamical transition temperature. The dynamical transition of protein has garnered various explanations. One theory suggests it’s due to the behavior of water in the hydration shell, transitioning from rigid to fluid at certain temperatures, thus influencing protein flexibility (7,12,16-19). Another theory considers the transition as an inherent property of the protein, where thermal energy allows the protein to access a wider range of conformations (21).

A prevailing scenario is that the internal dynamics of the protein is slaved to the motion of the surrounding hydration water (7,12,16-19), and thus the protein dynamical transition results from the changes in the dynamics of the hydration water with temperature (5,7,12,17,20). This scenario is indirectly supported by the experimental finding that the presence of the protein dynamical transition requires a minimum amount of hydration water, ∼ 0.2 gram water/gram protein (4,8). Further support comes from the results of all-atom molecular dynamics simulations, suggesting that it is the activation of the translational motions of surface water molecules around 200 K that leads to the dynamical transition in the underlying protein (5,7,20).

This ‘slaving’ scenario can be examined directly by an experiment using isotopic labelling in combination with elastic neutron scattering methods (7,21). Neutrons are highly sensitive to hydrogen atoms as their incoherent scattering cross section is an order of magnitude higher than the incoherent and coherent scattering cross sections of other elements (22-24). Thus, neutron signals collected on an ordinary protein powder hydrated in D_2_O reflect the dynamics of the protein while signals from the perdeuterated sample in H_2_O inform about the motion of water. The experimental results derived from this combined approach are, however, inconsistent (7,21,25). Measurements performed on perdeuterated maltose binding protein hydrated in H_2_O revealed a harmonic-to-anharmonic transition for hydration water taking place at the same temperature as that of the underlying protein (7). In contrast, a similar experiment on perdeuterated green fluorescence protein showed that the anharmonic onset in hydration water occurs at a lower temperature than that of the protein (21). More recent measurements on lysozyme hydrated in both D_2_O and H_2_O found that the transition temperature of protein and water coincided when examining their atomic displacements at 1 ns, but took place at different temperatures when changing the explored time scale to 3 ns (25). Therefore, there remains an unanswered question concerning whether the transition in dynamics of protein around 200 K is indeed coupled to that of the hydration water, whose resolution is of fundamental importance to understand the mechanism governing the nature of their interaction.

To address this, it requires a systematic measurement of the temperature dependence of atomic displacements of the protein and its surface water separately, as a function of hydration levels, *h* (gram water/gram protein), and at different time scales (instrument resolutions). Here, we performed elastic neutron scattering experiments on a number of protein powders hydrated in D_2_O and on the perdeuterated counterparts hydrated in H_2_O, to track the dynamics of protein and hydration water independently. Moreover, using a range of neutron instruments with distinct resolutions, we tested the effect of the explored time scales on the dynamics of the two components. Four globular proteins with different secondary and tertiary structures (see Fig.S1 and Table S1) were studied here. We found that the onset temperature (*T*_on_) of the protein dynamical transition varies with both biomolecular structure and hydration level, but is independent of the instrumental resolution time. Conversely, *T*_on_ of the hydration water is insensitive to both the protein structure and the level of hydration, but solely determined by the instrument resolution. Therefore, the dynamical transition of the protein is decoupled from the onset of anharmonic dynamics of its hydration water around 200 K. The onset in water cannot be assigned to a physical transition, but to a resolution effect. In contrast, the protein dynamical transition is an intrinsic change in the dynamics of the biomolecules. Complementary differential scanning calorimetry (DSC) measurements revealed a step-like change in the heat flow around the transition temperature of the protein, similar to the glass transition observed in polymers. This suggests that the dynamical transition in the protein results from a similar process involving the freezing of the structural relaxation of the protein molecules beyond equilibrium.

## Results

### Elastic Neutron Scattering Experiments

The quantity measured in the neutron experiment is the elastic intensity, i.e. the intensity of the elastic peak in the dynamic structure factor, *S*(*q*, Δ*t*), where *q* is the scattering wave vector and Δ*t* is the resolution time of the neutron spectrometer. *S*(*q*, Δ*t*) is an estimate of the average amplitude of the atomic motions up to Δ*t* (11,22). Three neutron backscattering spectrometers were chosen to cover a wide range of timescales; HFBS at the NIST Center for Neutron Scattering, USA, DNA at the Materials and Life Science Experimental Facility at J-PARC in Japan, and OSIRIS at the ISIS Neutron and Muon Facility, UK. The instrumental energy resolutions are 1 µeV, 13 µeV, 25.4 µeV and 100 µeV, corresponding to timescales of ∼1 ns, ∼80 ps, ∼40 ps, and ∼10 ps, respectively. Four globular proteins were investigated, myoglobin (MYO), cytochrome P450 (CYP), lysozyme (LYS), and green fluorescent protein (GFP), the detailed structural features of which are presented in Fig. S1 and Table S1. For simplicity, the hydrogenated and perdeuterated proteins are noted as H-protein and D-protein, respectively. Details of the sample preparation and neutron experiments are provided in Supplemental Material (SM).

### Dynamics of protein

Figs. 1a to g show the temperature dependence of *S*(*q*, Δ*t*) collected on H-LYS, H-MYO and H-CYP in dry and hydrated state with D_2_O measured by neutron spectrometers of different resolutions, Δ*t*. Since the measurements were performed on H-protein in D_2_O, the signals reflect the dynamics of the proteins. A clear deviation can be seen in the temperature dependence of *S*(*q*, Δ*t*) for the hydrated protein from that of the dry powder, which is defined as the onset temperature, *T*_on_ (8,12,25-27), of the protein dynamical transition. The advantage of such definition is that it highlights the effect of hydration on the anharmonic dynamics in proteins while removing the contribution from the local side groups, e.g., methyl groups, whose motions are hydration independent (11,28). As shown in Ref. (12,29), the activation temperature of the rotations of methyl group varies with the instrument resolution, which will cloud the present analysis. Two important conclusions can be drawn from Fig.1 (*T*_on_ is summarized in Table S5). (i) *T*_on_ is distinct for each protein, LYS (213 K), MYO (198 K) and CYP (228 K), and (ii) it is independent of the timescale explored even though the resolutions of the neutron spectrometers differ by orders of magnitude. Using the same set of data, we also analyzed the temperature dependence of the mean squared atomic displacements, <x^2^(Δt)> (see details in SM and results in Fig. S2) and obtained similar conclusions. We further calculated *T*_on_ of H-protein in D_2_O in the *q*-range from 0.45-0.9 Å ^− 1^ and 1.1-1.75 Å ^− 1^. As shown in Table S3-S4, the *q*-range does not alter the *T*_on_ of proteins.

**Figure 1.**
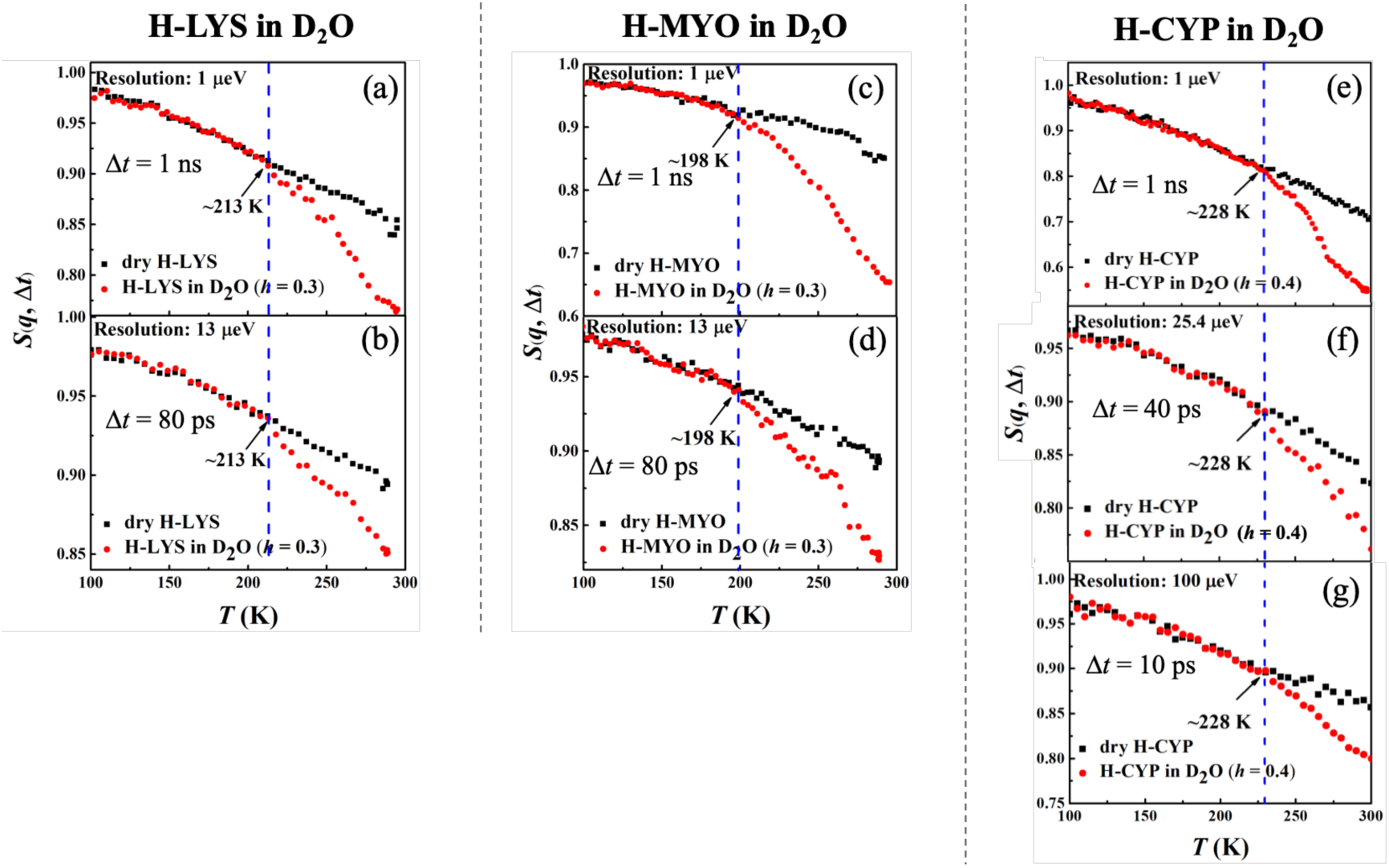
Resolution dependence of the onset of protein dynamical transition. Neutron spectrometers with different resolutions (1, 13, 25.4 and 100 µeV) were applied. Elatic intensity *S*(*q*, Δ*t*) of (a, b) dry H-LYS and H-LYS in D_2_O at *h* = 0.3, (c, d) dry H-MYO and H-MYO in D_2_O at *h* = 0.3, and (e-g) dry H-CYP and H-CYP in D_2_O at *h* = 0.4. All the experimental *S*(*q*, Δ*t*) are normalized to data measured at ∼10 K and summed over values of *q* ranging from 0.45 to 1.75 Å^−1^. The dashed lines in each figure identify the onset temperatures of the transition, *T*_on_, where the neutron data of the hydrated system deviate from the dry form. The same analyses are used in Figs. 2 to 4 and S2 to S5.

These findings suggest that the dynamical transition in the protein is an intrinsic property of the hydrated biomolecule, and it depends on the structure and chemistry of the protein concerned. Our results are consistent with Ref. (12), which demonstrated that *T*_on_ in both protein and polypeptide is independent of the resolution of the neutron spectrometer, if one carefully removes the contributions from methyl rotations and vibrations to <x^2^(Δt)> measured by elastic neutron scattering. Additionally, Ref. (25) showed that, as compared to the dry form, the D_2_O-hydrated lysozyme presents an approximately resolution-independent *T*_on_, again in agreement with our findings.

Fig. 2 compares the temperature dependence of *S*(*q*, Δ*t*) measured on H-CYP and H-LYS in D_2_O at different hydration levels, *h* (*T*_on_ is summarized in Table S6). Evidently *T*_on_ of the protein increases from 228 to 248 K when reducing *h* from 0.4 to 0.2 (Figs. 2a) by using the neutron instrument with Δ*t* = 1 ns. A similar hydration dependence of *T*_on_ is also observed when we replot the neutron data measured on H-LYS hydrated in D_2_O reported in Ref. (26) (Fig. 2c). It can be found that *T*_on_ of lysozyme changes systematically from 195 to 225 K, when decreasing *h* from 0.45 to 0.18. Similar conclusion can be obtained when we analyzed < x^2^(Δt)> (see Fig. S3). The dynamical transition temperature in lipid membranes is higher when the membrane is dry (51). We also studied the secondary structure content and tertiary structure of CYP protein at different hydration levels (*h* = 0.2 and 0.4) through molecular dynamics simulation. As shown in Table S2 and Fig. S6, the extent of hydration does not alter the protein secondary structure content and overall packing. Thus, this result suggests that water molecules have more influence on protein dynamics than on protein structure.

**Figure 2.**
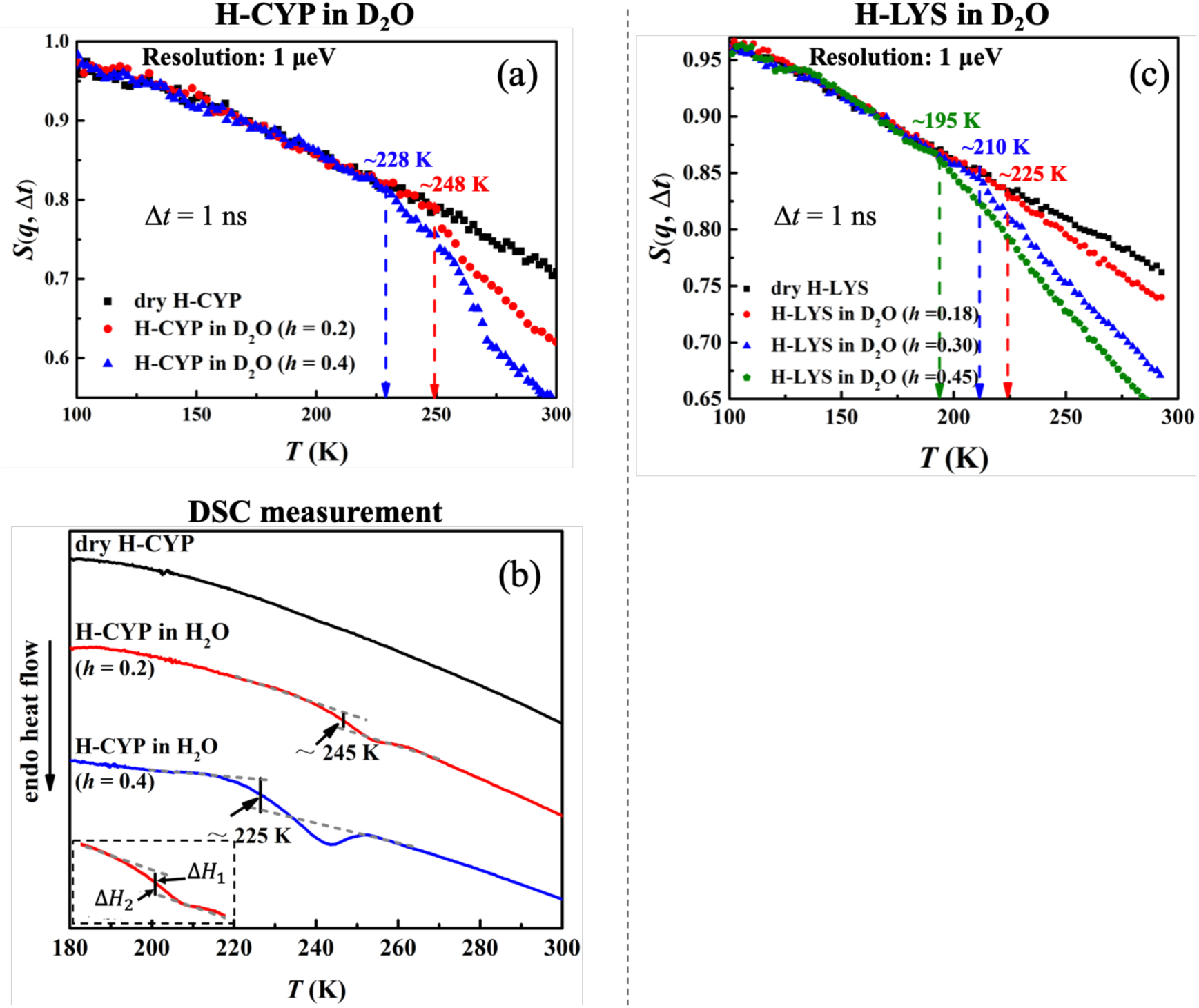
Hydration dependence of the onset of protein dynamical transition. *S*(*q*, Δ*t*) of (a) dry H-CYP and H-CYP in D_2_O at *h* = 0.2 and 0.4 and (c) dry H-LYS and H-LYS in D_2_O at *h* = 0.18, 0.30 and 0.45, all measured using HFBS with the instrumental resolution of 1 μeV. All the Data in (c) were replotted from Ref. (26). (b) DSC curves obtained for dry H-CYP and H-CYP in water at *h* = 0.2 and 0.4. *T*_DSC_ is defined as the midpoint between two heat flow baselines, where (35,36,49).

The results from the neutron scattering experiments suggest that the dynamical transition in proteins is an intrinsic property of the biomolecule and strongly depends on the amount of water surrounding it. Such an intrinsic transition can result either from a critical phase transition, e.g., water to ice (30,31), or from freezing of the structural relaxation of the system beyond the equilibrium time (∼100 - 1000 s) of the experiment, in analogy to the glass transition in polymers from rubbery state to the glass form (32-34). Both of them will significantly increase the mechanical modulus of the material and suppress the atomic displacements at the fast time scales (pico-to-nanosecond) probed by the neutron spectrometers (30-34) like those used in this work. To explore the microscopic nature of the protein dynamical transition, we performed differential scanning calorimetry (DSC) measurements on CYP at dry, *h* = 0.2 and 0.4. As illustrated by Fig. 2b, H-CYP at *h* = 0.2 and *h* = 0.4 exhibit a step-like transition in the heat flow at 245 K and 225 K, respectively, while no such transition is observed in dry H-CYP. Such step-like transition in heat flow is normally defined as the glass transition in polymers (35,36).

For simplicity, the step-like transition identified by DSC is noted as *T*_DSC_. When comparing Figs 2b with Fig. 2a, one can find that the values of *T*_DSC_ approximate those of *T*_on_ probed by neutrons. *T*_DSC_ of hydrated myoglobin was reported by literature to be 190 K (37), which is again in good agreement with the value of *T*_on_ in Fig. 1. More importantly, *T*_DSC_ and *T*_on_ present the same hydration dependence, i.e., both increase with decrease of *h* (see Figs. 2a and 2b). Therefore, we can conclude that the onset of anharmonicity around 200 K in proteins measured by neutron scattering as shown in Figs. 1 and 2 results from the freezing of the structural relaxation of the biomolecule beyond the equilibrium when cooling the system below *T*_DSC_, similar to the glass transition in polymers. Similar interpretation has also been suggested in Ref.(38).

As the time scale probed by neutron spans from pico-to nanoseconds, it is too fast to allow us to directly “see” structural relaxations of the protein around *T*_on_. However, “freezing” of the structural relaxation beyond the equilibrium time (∼100 - 1000 s), i.e., the measurement time of neutron experiments at each temperature, will turn the system into a “frozen” solid form, which can significantly suppress the fast dynamics measured by neutron and cause the transition probed (32-34,38). Moreover, water can be considered here as lubricant or plasticizer which facilitates the motion of the biomolecule (28,39,40). As widely observed in polymeric systems (41-43), adding water as plasticizer will significantly reduce the glass transition temperature of the polymers. This rationalizes the hydration effect on *T*_DSC_ and *T*_on_, both decreasing with increase of *h*.

### Dynamics in hydration water

Figs. 3a-b and c-e show the temperature dependence of *S*(*q*, Δ*t*) measured on perdeuterated GFP and CYP in dry and H_2_O-hydrated forms. In these samples, the measured signal reflects primarily the motions of water molecules. Two important observations arise from the data. Firstly, *T*_on_ of hydration water for these two proteins strongly depends on the resolution of the spectrometer, increasing drastically from 200 K to 250 K when reducing Δ*t* from 1 ns to 10 ps. Similar conclusions are obtained when we analyzed the temperature dependence of <x^2^(Δt)> (see Fig. S4). The observation of a dependence of *T*_on_ on Δ*t* is typical for a thermally activated process, which occurs when the characteristic relaxation time becomes comparable to the instrumental resolution, and the relaxation process is said to enter the time window of the instrument (29,44). In this case, it means that the relaxation time, *τ*, of hydration water is 10 ps at 250 K, 40 ps at 234 K and 1 ns at 200 K. Assuming an Arrhenius-type process, 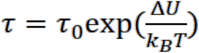, the energy barrier *ΔU* can be estimated to be ∼38 kJ/mol. Our results thus demonstrate that the anharmonic onset of the hydration water is in fact not a real physical transition but merely a resolution effect. It occurs as the relaxation time *τ* of water, which varies continuously with temperature, happens to cross the instrument resolution, Δ*t*, on the pico-to-nanosecond time scales at *T*_on_. Our findings agree with reports from dielectric measurements, the signal of which is highly sensitive to the rotation of hydration water (45,46). They showed a smooth temperature dependence of the characteristic relaxation time in the range from 170 K to 250 K without any sudden changes (45,46). Moreover, our data also agree with Ref. (47), which demonstrated that the characteristic relaxation time of protein-surface water, measured on H_2_O-hydrated perdeuterated C-phycocyanin by quasi-elastic neutron scattering, changes smoothly over temperature without any disruptions around the dynamical transition temperature of the protein. Secondly, the onset temperature of the hydration water is independent of the protein structure when Δ*t* is fixed, since the values of both GFP and CYP are identical.

**Figure 3.**
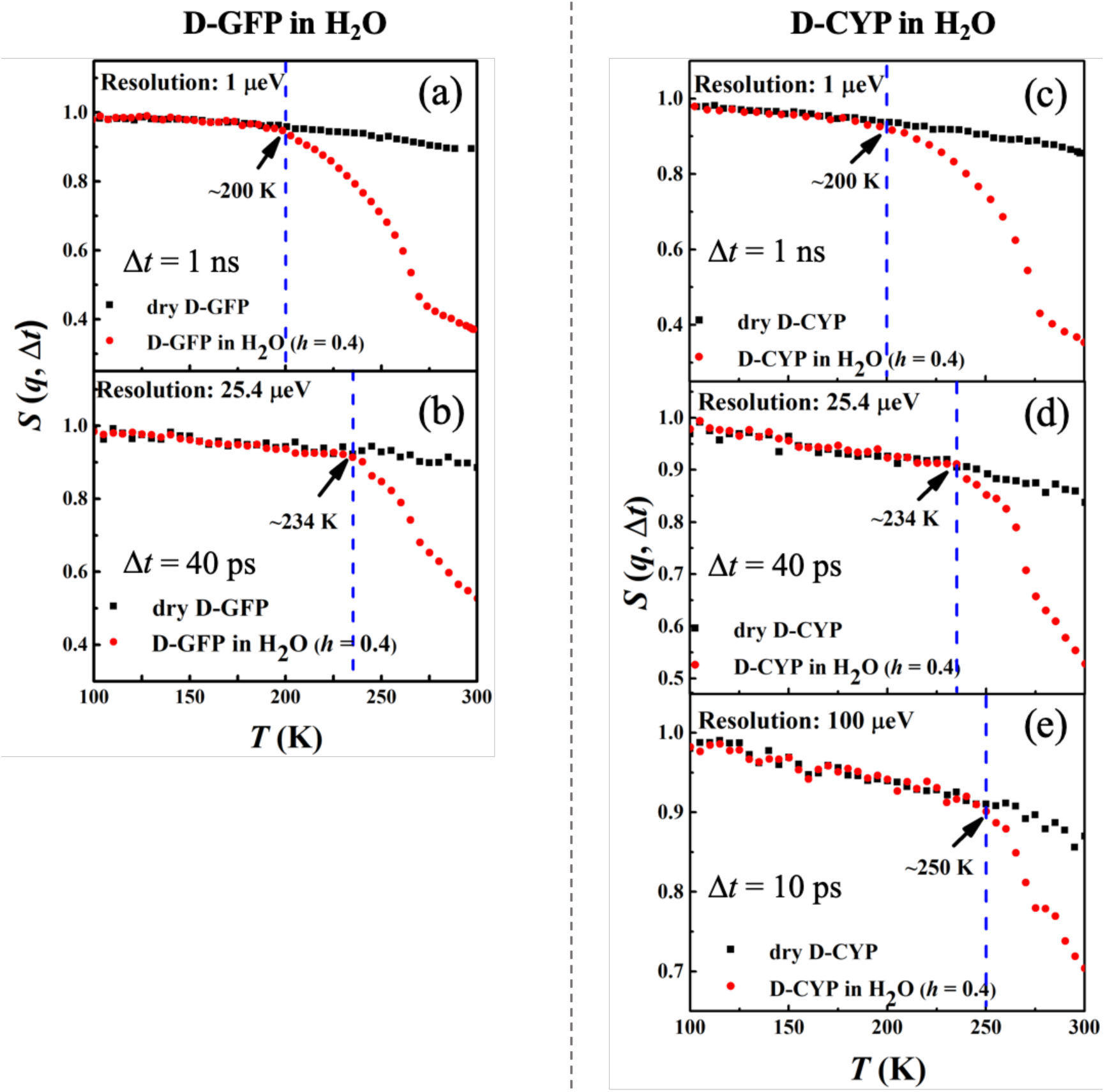
Resolution dependence of the anharmonic onset of hydration water. Neutron spectrometers with different resolutions (1, 25.4 and 100 µeV) were applied. *S*(*q*, Δ*t*) of (a, b) dry D-GFP and D-GFP in H_2_O at *h* = 0.4, and (c-e) dry D-CYP and D-CYP in H_2_O at *h* = 0.4.

Furthermore, the hydration dependence of the anharmonic onset of the water is presented in Fig. 4, which shows that *T*_on_ remains constant with *h* as long as the instrument resolution is fixed. This behavior is drastically different from that of the protein (Fig. 2). The same conclusions can be obtained when analyzing <x^2^(Δt)> (see Figs. S5).

**Figure 4.**
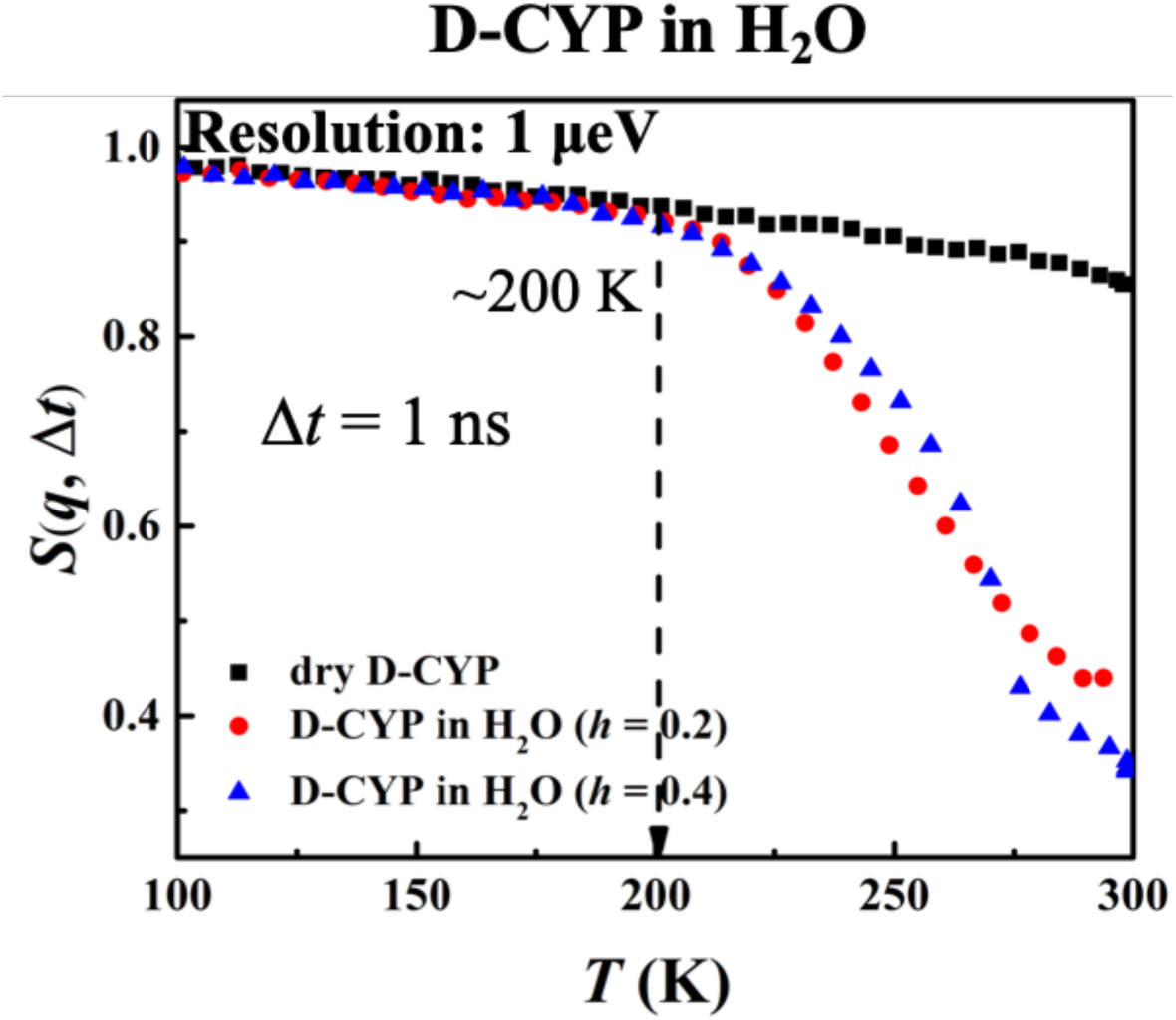
Hydration dependence of the anharmonic onset of hydration water. *S*(*q*, Δ*t*), for dry D-CYP and D-CYP in H_2_O at *h* = 0.2 and 0.4, measured using HFBS neutron instrument with an energy resolution of 1 µeV.

## Conclusion and Discussion

By combining elastic neutron scattering with isotopic labelling, we have been able to probe the dynamics of the protein and surface water separately, as a function of temperature, protein structural composition, hydration level and time scale. We found that the anharmonic onsets of the two components around 200 K are clearly decoupled and different in origin. The protein shows an intrinsic transition, whose temperature depends on the structure of the protein and the hydration level, and not on the instrument used to measure it. It has a thermodynamic signature similar to the glass transition in polymers as confirmed by DSC, and thus can be assigned to the freezing of the structural relaxation of the protein beyond the experimental equilibrium time (100 to 1000 s). In contrast, the temperature at which the onset of anharmonicity happens in the hydration water is given by the instrument resolution, independent of both the biomolecular structure and the level of hydration.

Based on our findings, we can infer that, in some cases, the dynamical transition of a protein can coincide with the anharmonic onset of its surface water if one characterizes the system using a single neutron instrument with a fixed resolution. But such coincidence will be torn apart if the measurements were conducted by using instruments of different resolutions or at different amounts of hydration, such as in the present work. This rationalizes the seemingly contradictive results reported in the literature (7,21,25).

The protein dynamical transition has long been thought to connect to the thermal onset of the functionality of the biomolecule. Our experiments suggest that this transition in protein is an intrinsic property of the hydrated protein that its structural relaxation is activated upon heating above the onset temperature. This structural relaxation might be associated with conformational jumps of the biomolecules among different functional states, such as the states with the ligand-binding pocket being opened or closed. Unfreezing of the protein structural relaxation might facilitate these conformational jumps, turning on its functionality. However, as revealed by Ref (48), the denatured form of lysozyme also exhibits a dynamical transition, similar to that seen in its folded native form. Additionally, the dynamical transition also can be found in the mixture of amino acids (12). Hence, one can argue that the activation of the structural relaxation of the biomolecule above the dynamical transition temperature is a necessary but insufficient condition for the protein to function, as the latter also requires the biomolecule assuming the correctly folded 3-dimensional structure. The findings in this work help further the understanding of the microscopic mechanism governing the dynamics in proteins and their hydration water, as well as their interactions at the cryogenic temperature. More importantly, we demonstrated that the protein dynamical transition is a real transition, connecting to unfreezing of the biomolecular structural relaxation, which could be crucial for activating the function.

## Acknowledgments

This work is supported by the National Natural Science Foundation of China (11974239; 62302291), the Innovation Program of Shanghai Municipal Education Commission (2019-01-07-00-02-E00076). H.O’N. and Q.Z. acknowledge the support of Center for Structural Molecular Biology (FWPERKP291) funded by the U.S. Department of Energy (DOE) Office of Biological and Environmental Research. Access to the HFBS was provided by the Center for High-Resolution Neutron Scattering, a partnership between the National Institute of Standards and Technology and the National Science Foundation under Agreement No. DMR-1508249. The neutron experiment at the Materials and Life Science Experimental Facility of the J-PARC was performed under a user program (Proposal No. 2019A0020). We thank STFC for access to neutron scattering facilities at RB1800112. The original data is accessible via data cite: htpps://doi.10.5286/ISIS.E.RB1800112.

