## Supplementary material for "Decoupling of the Onset of Anharmonicity between a Protein and Its Surface Water around 200 K": SI

Context:
Materials and Methods
Table S1-S6 and Figure S1-S7
References

**Sample preparation**

Hydrogenated LYS from chicken egg white and hydrogenated MYO from equine skeletal muscle were purchased from Sigma Aldrich (Shanghai, China). The expression and purification of hydrogenated and perdeuterated CYP (We used P450 from Pseudomonas putida for the study) and GFP are described previously [1-4]. In order to exclude the effect of ions, the proteins were dialyzed before experiments. For simplification, the hydrogenated protein and perdeuterated protein are denoted as H-proteins and D-proteins in the manuscript, respectively. All the H-proteins were dissolved in $D_2O$ to allow full deuterium exchange of all exchangeable hydrogen atoms and then lyophilized for 12 hours to obtain the dry sample. The lyophilized H-protein is then put into a desiccator with $D_2O$, placed in the glove box purged with nitrogen gas, to absorb $D_2O$ till the desired hydration level, $h$ (gram water/gram protein). In contrast, the preparation of the deuterated proteins was conducted in the opposite way. The D-proteins were dissolved in $H_2O$ to allow full hydrogen exchange of all exchangeable deuterium atoms and then lyophilized for 12 hours to obtain the dry sample. The lyophilized D-protein is then put into a desiccator with $H_2O$ to absorb $H_2O$ till the desired $h$. The ultrapure water ($H_2O$) was supplied by a Millipore Direct-Q system (18.2 MΩ·cm at 25 °C). The deuterium oxidized ($D_2O$, 99.9 atom % D) was purchased from Sigma-Aldrich (Shanghai, China). The hydration levels of protein samples were controlled by measuring the sample weights before and after water adsorption. In this work, $h$ ranges from 0.02 (lyophilized dry form), 0.2, 0.3 to 0.4, where $h$ = 0.4 corresponds to a case that roughly a single layer of water molecules covers the protein's surface [5,6]. The dry H-CYP, H-LYS, H-MYO, and their

$D_2O$-hydrated forms at $h$ = 0.2, 0.3 or 0.4, and the dry D-GFP and D-CYP, and their $H_2O$-hydrated powders at $h$ = 0.4 are prepared for neutron scattering experiments. The accuracy of $h$ is controlled within 10% error. E.g., $h$ = 0.4±0.04 gram water/gram protein. All samples were sealed tightly in the aluminum cans in nitrogen before the neutron scattering experiments.

The dry H-CYP lyophilized in $H_2O$ and the ones hydrated in $H_2O$ at $h$ = 0.2 and 0.4 are prepared for the differential scanning calorimetry (DSC) measurement.

**Elastic incoherent neutron scattering (EINS)**

The elastic scattering intensity $S(\boldsymbol{q}, \Delta t) \approx I_{inc}(\boldsymbol{q}, \Delta t) = \frac{1}{N}\sum_j^N b_{j,inc}^2 \langle \exp[-i\boldsymbol{q}\cdot\boldsymbol{r}_j(0)]\exp[i\boldsymbol{q}\cdot\boldsymbol{r}_j(\Delta t)]\rangle$ is normalized to the lowest temperature (~10 K) and is approximately the value of the intermediate scattering function when decaying to the instrument resolution time, $\Delta t$ [1]. All the $S(q, \Delta t)$ was obtained in the temperature range of ~10 to 300 K during heating process with the rate of 1.0 K/min by using the HFBS at NIST, DNA at J-PARC and OSIRIS at ISIS. The energy resolutions of HFBS, DNA and OSIRIS are 1 μeV, 13 μeV, 25.4 μeV and 100 μeV, corresponding to the resolution times of ~1 ns [7], ~80 ps [8,9], ~40 ps [10] and ~10 ps [10], respectively. The results from instruments with various resolutions were summed over the same $q$ from 0.45 to 1.75 $Å^{-1}$.

**Differential scanning calorimetry (DSC).**

DSC measurements were performed by using the METTLER instruments DSC3+. The sample was sealed in a pan of aluminum. An empty pan was used as a reference. All the experiments were carried out in the temperature ranged from 150 to 300 K with heating rate of 1 K/min. The heating rate of DSC is the same as neutron experiments.

**Estimation of the Mean-squared atomic displacement (MSD).**

The mean-squared atomic displacement $< x^2(\Delta t) >$ was estimated by performing Gaussian approximation, where $S(q, \Delta t) = \exp(-\frac{1}{6}q^2 < x^2(\Delta t) >$. The values of $q$ used for Gaussian fitting ranges from 0.45 to 0.9 $Å^{-1}$ [11].

**Molecular dynamics simulation.**

The initial structure of protein cytochrome P450 (CYP) for simulations was taken from PDB crystal structure (2ZAX). Two protein monomers were filled in a cubic box. 1013 and 2025 water molecules were inserted into the box randomly to reach a mass ratio of 0.2 and 0.4 gram water/1 gram protein, respectively, which mimics the experimental condition. Then 34 sodium counter ions were added to keep the system neutral in charge. The CHARMM 27 force field in the GROMACS package was used for CYP, whereas the TIP4P/Ew model was chosen for water. The simulations were carried out at a broad range of temperatures from 360 K to 100 K, with a step of 5 K. At each temperature, after the 5000 steps energy-minimization procedure, a 10 ns NVT is conducted. After that, a 30 ns NPT simulation was carried out at 1 atm with the proper periodic boundary condition. As shown in Fig. S7, 30 ns is sufficient to equilibrate the system. The temperature and pressure of the system is controlled by the velocity rescaling method and the method by Parrinello and Rahman, respectively. All bonds of water in all the simulations were constrained with the LINCS algorithm to maintain their equilibration length. In all the simulations, the system was propagated using the leap-frog integration algorithm with a time step of 2 fs. The electrostatic interactions were calculated using the Particle Mesh Ewalds (PME) method. A non-bond pair-list cutoff of 1 nm was used and the pair-list was updated every 20 fs. All MD simulations were performed using GROMACS 4.5.1 software packages.

**Protein samples used for experiments:**

We studied four globular proteins, myoglobin (MYO), cytochrome P450 (CYP), lysozyme (LYS), and green fluorescent protein (GFP), the detailed structural features of which are presented in Fig. S1 and Table S1. The four proteins differ significantly in both secondary and tertiary structures. MYO is primarily a helix protein while GFP is dominated by beta sheets. Moreover, LYS contains two structural domains linked by a hinge while the other three are single-domain proteins.

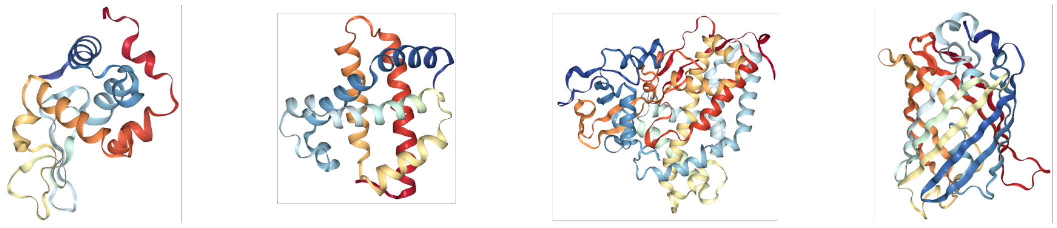

(a) Lysozyme (b) Myoglobin (c) Cytochrome P450 (d) Green fluorescent protein

Figure S1. Structures of (a) lysozyme (LYS), (b) myoglobin (MYO), (c) cytochrome P450 (CYP) and (d) green fluorescent protein (GFP).

Table S1. Relative content of each secondary structure in the proteins

| **Protein** | **Lysozyme** | **Myoglobin** | **Cytochrome P450** | **Green Fluorescent Protein** |
|---|---|---|---|---|
| Abbreviation | LYS | MYO | CYP | GFP |
| PDB ID | 1AKI | 2V1K | 2ZAX | 1EMB |
| $\alpha$-Helix* | 40% | 76% | 52% | 7% |
| $\beta$-Sheet* | 12% | 0% | 11% | 50% |
| Loop and Turn* | 48% | 24% | 37% | 43% |

*The relative content of each secondary structure is defined by mass fraction.

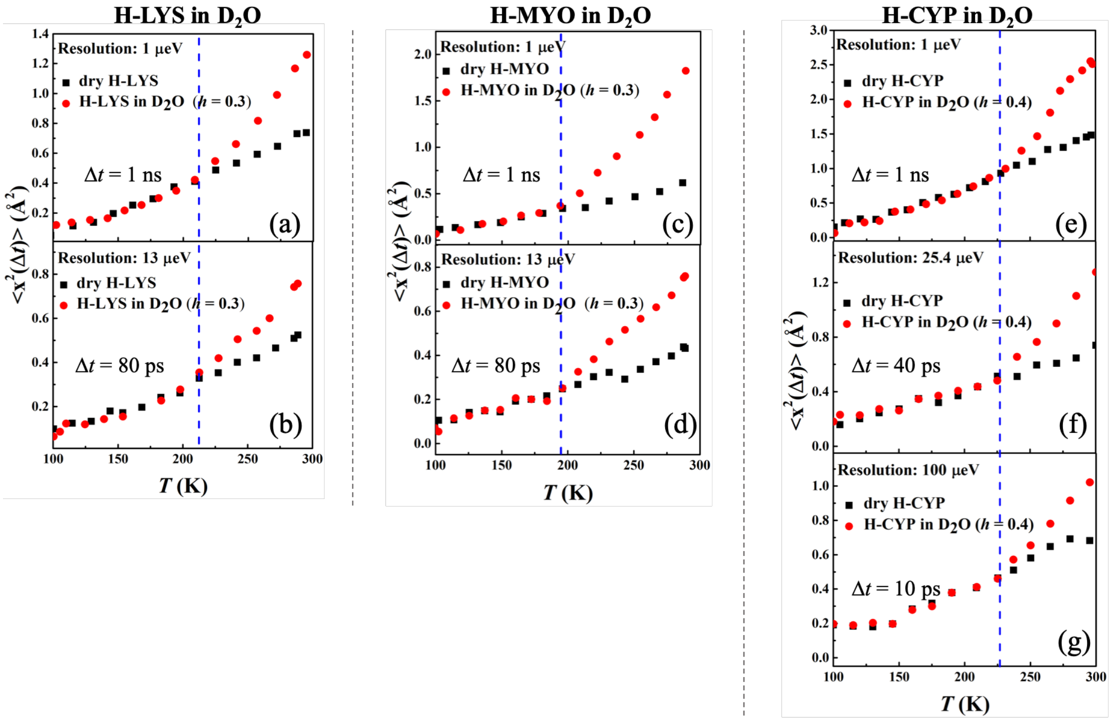


Figure S2. Resolution dependence of the onset of protein dynamical transition. Mean-squared atomic displacements $< x^2(\Delta t) >$, derived from Figure 1 using Gaussian approximation, of (a, b) dry H-LYS and H-LYS in $D_2O$ at $h$ = 0.3, (c, d) dry H-MYO and H-MYO in $D_2O$ at $h$ = 0.3, and (e-g) dry H-CYP and H-CYP in $D_2O$ at $h$ = 0.4.

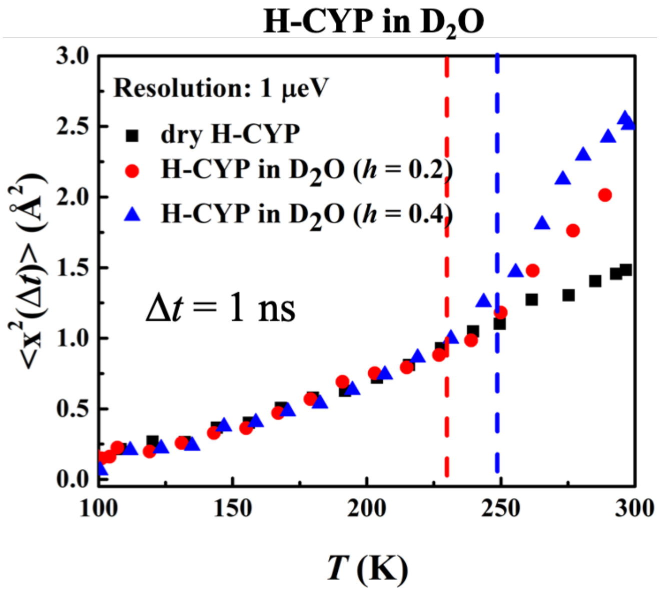


Figure S3. Resolution dependence of the onset of protein dynamical transition. $< x^2(\Delta t) >$, derived from Figure 2(a) using Gaussian approximation, of dry H-CYP and H-CYP in $D_2O$ at $h$ = 0.2 and 0.4.

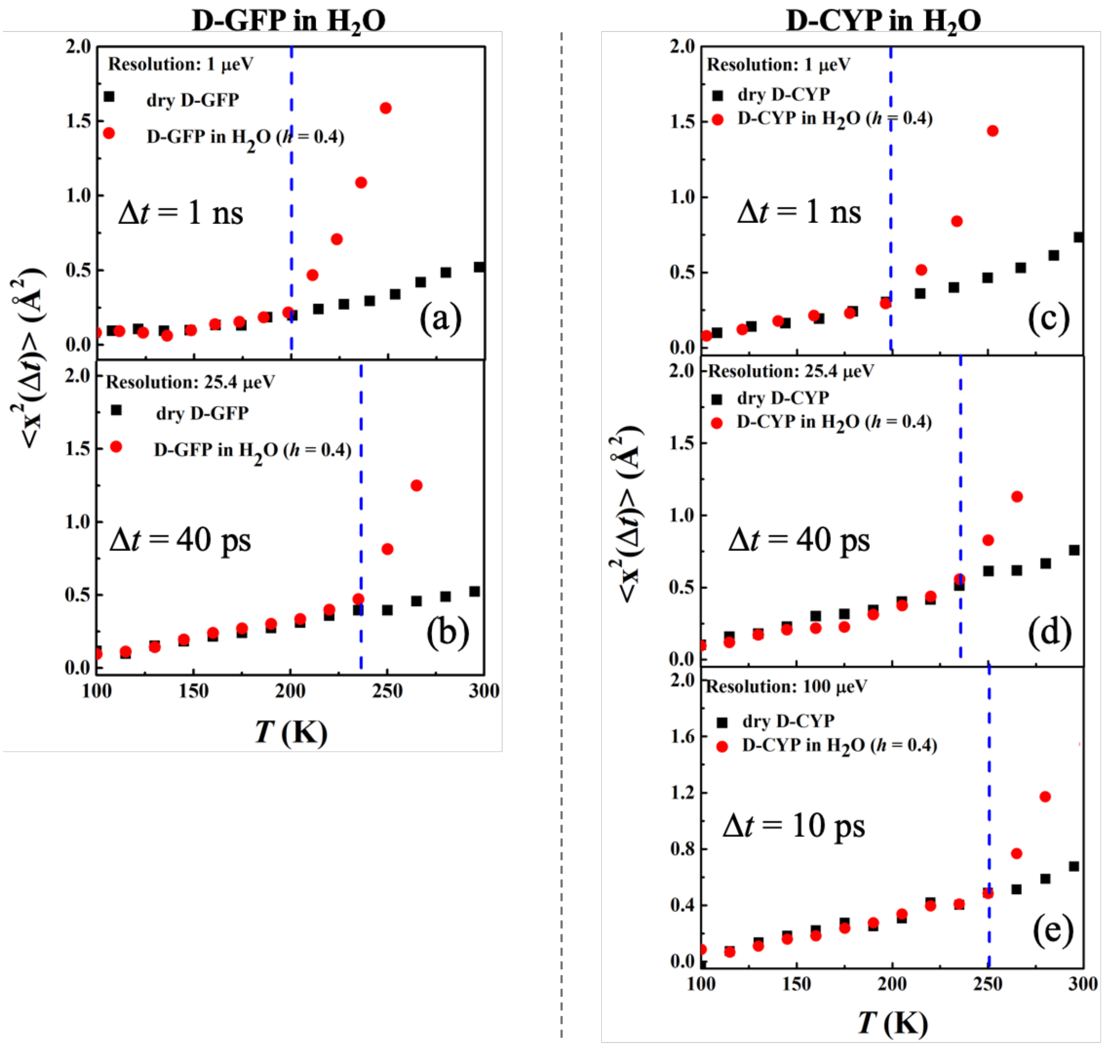


Figure S4. Resolution dependence of the anharmonic onset of hydration water. Mean-squared atomic displacements $< x^2(\Delta t) >$, derived from Figure 3 using Gaussian approximation, of (a, b) dry D-GFP and D-GFP in $H_2O$ at $h$ = 0.4, (c-e) dry D-CYP and D-CYP in $H_2O$ at $h$ = 0.4.

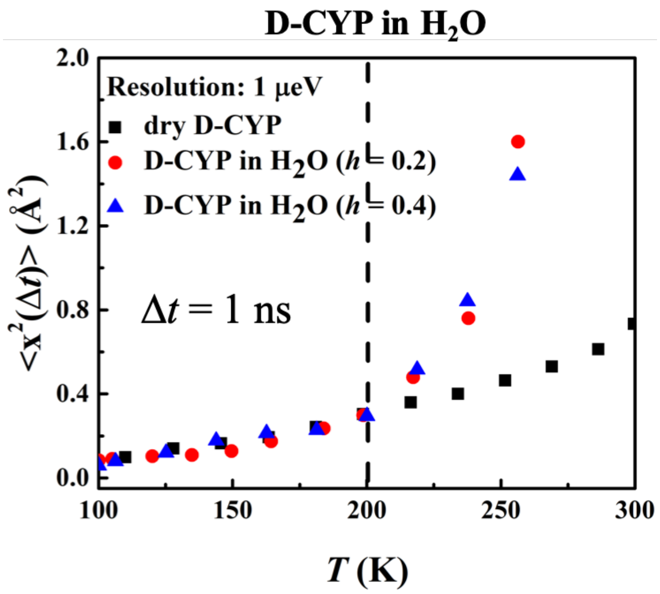


Figure S5. Hydration dependence of the anharmonic onset of hydration water. $< x^2(\Delta t) >$, derived from Figure 4 using Gaussian approximation, of dry D-CYP and D-CYP in $H_2O$ at $h$ = 0.2 and 0.4.

Table S2. The secondary structure content of CYP protein at different hydration levels

| | alpha-helix | beta-sheet | loop and turn |
|---|---|---|---|
| CYP ($h$ = 0.2) | 52% | 11% | 37% |
| CYP ($h$ = 0.4) | 52% | 11% | 37% |

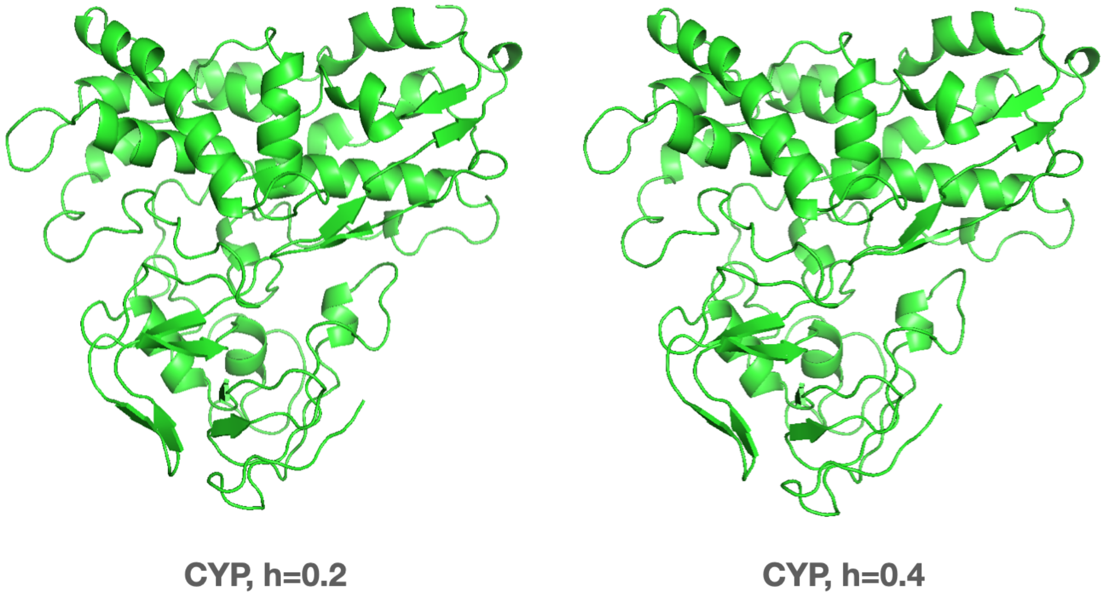


Figure S6. The 3D structure of CYP protein at different hydration levels obtained from MD simulations (PDB ID: 2ZAX).

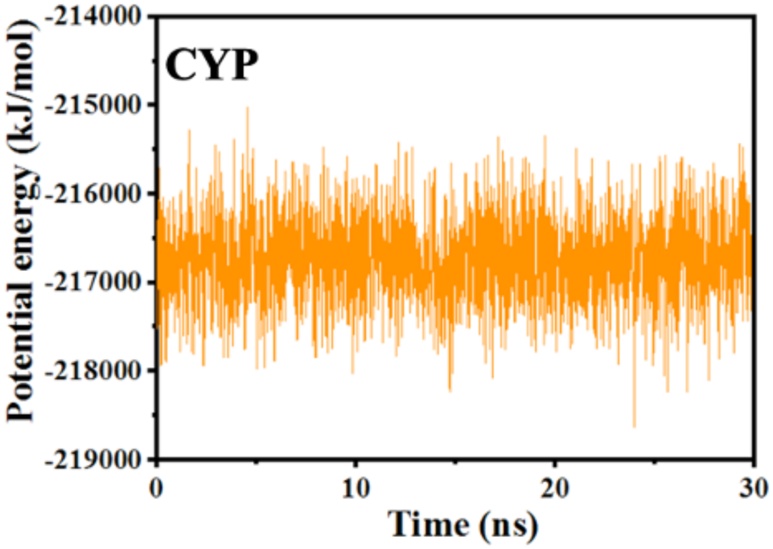


Figure S7. The potential energy as a function of MD trajectory time of CYP.

Table S3. $T_{on}$ of protein in q-ranges from q = 0.45-0.9 $Å^{-1}$

| | 1 ns | 80 ps | 40 ps | 10 ps |
|---|---|---|---|---|
| LYS | 213 K | 213 K | - | - |
| MYO | 198 K | 198 K | - | - |
| CYP | 228 K | - | 228 K | 228 K |

Table S4. $T_{on}$ of protein in q-ranges from q = 1.1-1.75 Å$^{-1}$

| | 1 ns | 80 ps | 40 ps | 10 ps |
|---|---|---|---|---|
| LYS | 212 K | 213 K | - | - |
| MYO | 197 K | 199 K | - | - |
| CYP | 228 K | - | 227 K | 228 K |

Table S5. $T_{on}$ of protein at different time resolution

| | 1 ns | 80 ps | 40 ps | 10 ps |
|---|---|---|---|---|
| LYS (h = 0.3) | 213 K | 213 K | - | - |
| MYO (h = 0.3) | 198 K | 198 K | - | - |
| CYP (h = 0.4) | 228 K | - | 228 K | 228 K |

Table S6. $T_{on}$ of protein at different hydration level

| | 0.18 | 0.2 | 0.3 | 0.4 | 0.45 |
|---|---|---|---|---|---|
| LYS (1 ns) | 225 K | - | 213 K | - | 195 K |
| CYP (1 ns) | - | 248 K | - | 228 K | - |
| CYP ($T_{DSC}$) | - | 245 K | - | 225 K | - |
